# Barcoding Notch signaling in the developing brain

**DOI:** 10.1101/2024.05.10.593533

**Authors:** Abigail Siniscalco, Roshan Priyarangana Perera, Jessie E. Greenslade, Aiden Masters, Hannah Doll, Bushra Raj

## Abstract

Developmental signaling inputs are fundamental for shaping cell fates and behavior. However, traditional fluorescent-based signaling reporters have limitations in scalability and molecular resolution of cell types. We present SABER-seq, a CRISPR-Cas molecular recorder that stores transient developmental signaling cues as permanent mutations in cellular genomes for deconstruction at later stages via single-cell transcriptomics. We applied SABER-seq to record Notch signaling in developing zebrafish brains. SABER-seq has two components: a signaling sensor and a barcode recorder. The sensor activates Cas9 in a Notch-dependent manner with inducible control while the recorder accumulates mutations that represent Notch activity in founder cells. We combine SABER-seq with an expanded juvenile brain atlas to define cell types whose fates are determined downstream of Notch signaling. We identified examples wherein Notch signaling may have differential impact on terminal cell fates. SABER-seq is a novel platform for rapid, scalable and high-resolution mapping of signaling activity during development.

## INTRODUCTION

Neural cell fate decisions are influenced by many factors including a cell’s division history (lineage), spatial location (niche) and the signaling inputs received^1^. Pioneering studies have shown that cellular signaling is an important mechanism for establishing the embryonic brain framework as it regulates patterning, morphogenesis, migration, proliferation and differentiation^2^. Examples of well-studied signaling pathways involved in neurogenesis include Notch, Fibroblast Growth Factor (Fgf), Sonic Hedgehog (Shh), Wnt, and Bone Morphogenetic Proteins (BMP)^3^. Classically, cellular signal transduction has been studied using signaling reporters that couple ligand or receptor activation with a downstream readout such as fluorescent gene expression or chemical luminescence. The advantages of such reporters include identification and quantification of cells that received signaling inputs, quantification of the signaling strength, and depending on the sensitivity of the reporter, measurement of signaling dynamics. However, studies utilizing these reporters are limited in two main aspects. First, imaging-based signaling readouts are limited in throughput as relatively small populations of cells localized to a predefined geographical area are visualized due to time and cost constraints. Second, these methods do not provide high resolution cell type identification, which is especially important in highly heterogenous tissues such as the brain.

Single-cell genomics has become the method of choice for characterizing molecular states and cell type identities with unprecedented throughput and fine resolution in a plethora of organisms and in vitro systems^4^. Recently, single-cell genomics has been coupled with innovative genetic editing methods to characterize various biological paradigms. For example, CRISPR-Cas genome editing tools and single-cell RNA sequencing (scRNA-seq) have been combinatorially utilized to perform clonal tracing and lineage tree reconstruction with cell type resolution in development and disease^5,6^. Many iterations of CRISPR-based lineage recorders, also referred to as molecular recorders, have been described. One group of recorders rely on generation of a diverse collection of edited barcodes via Cas9-induced deletions to transgenic arrays or endogenous loci, which are then decoded by sequencing^7–14^. Another group of recorders, known as “DNA writers”, are designed to predominantly generate small insertions to a barcode array and circumvent loss of information often observed with deletion-based Cas9 editors^15–22^. A third group utilizes orthogonal Cas proteins such as Cas12a, including nuclease-dead base editor versions, for multiplexed event recordings^23,24^. In addition to lineage analyses, studies have applied molecular recorders to record a wide array of cellular and synthetic events such as gene expression, cellular stimuli, synthetic signals, videos, and text messages but these have been limited to cell culture systems or prokaryotic cells and have not been combined with scRNA-seq ^15–22^. However, biological applications of molecular recorders using in vivo animal models have been restricted to cell lineage and clonal analyses.

We reasoned that CRISPR-Cas molecular recorders can be adapted to investigate a new paradigm - signaling inputs during animal development. Here we describe a new method, SABER (<u>s</u>ignal-<u>a</u>ctivated <u>b</u>arcode <u>e</u>diting recorder), where signaling-induced activation of Cas9 promotes in vivo barcode editing. We apply SABER to capture Notch signaling during brain organogenesis in zebrafish. By coupling SABER to scRNA-seq (SABER-seq) we identify brain cell types that were derived from Notch-stimulated progenitor cells. Compared to traditional signaling reporters, SABER enhances cell type resolution, and enables permanent genomic recording of transient signaling events to be recovered at later timepoints. The latter provides a means to assess the impact of signal transduction on cell fate decisions. SABER has been designed to be modular, and thus the various genetic components can be swapped to study any signaling event at any timepoint in any organism or in vitro system of interest.

## RESULTS

### TP1 enhancer driven Notch signal recorder

A simple design for a signal-activated CRISPR-Cas recorder is to induce Cas9 expression in a signal-dependent manner, and subsequently record this signaling activity via Cas9-mediated mutations (edits) to a genomic barcode array. To test this idea, we focused on the Notch signaling pathway since it is fundamental for many aspects of development, including neurogenesis where one of its functions is to promote progenitor cell maintenance and prevent precocious neuronal differentiation^25,26^. Furthermore, since it is a well-characterized pathway, numerous Notch-responsive cis-elements and small molecule modulators of Notch signaling have been described in zebrafish, which we leveraged in the design of our study. The TP1 enhancer element, comprising 12 copies of Notch co-factor binding sites (signaling response element) upstream of a minimal promoter has been shown to be Notch responsive in mammals and zebrafish^27^. Moreover, it is active in Notch-responsive tissues such as the central nervous system, pancreas, vasculature, liver and intestine^27^. We generated transgenic zebrafish with a TP1 enhancer driving Cas9-GFP expression and U6 promoters driving constitutive zygotic expression of four sgRNAs (TP1:Cas9-GFP, 4[U6:sg]). Comparison of TP1:GFP and TP1:Cas9-GFP, 4[U6:sg] transgenic lines revealed similar GFP expression patterns with fluorescence detected in the forebrain, hindbrain, spinal cord, and vasculature, among other regions (Fig. 1a, b). However, we observed that TP1:Cas9-GFP, 4[U6:sg] transgenics had weaker GFP expression likely due to the large size of the transgenic construct, and GFP expression was barely detectable beyond 48-72 hours post fertilization (hpf). We observed similar Cas9-GFP silencing in other transgenic lines (e.g. b-actin promoter [Olactb] driven Cas9-GFP, data not shown). This suggested that widespread and constitutive in vivo expression of Cas9 may not be well tolerated in developing zebrafish. To rapidly screen our transgenics for functional Cas9 activation and editing capacity, we designed one of the sgRNAs to target the tyrosinase gene, *tyr*, which is necessary for pigment formation. We outcrossed TP1:Cas9-GFP, 4[U6:sg] founders to wildtype zebrafish and observed that F1 adults had prominent stripe mutations in mosaic patterns (Fig 1c). Indeed, Notch signaling has been implicated in melanophore survival and adult zebrafish stripe patterning^28^. F1 adults generated from outcrossing Olactb:Cas9-GFP, 4[U6:sg] founders had more extensive pigmentation disruption (including near complete loss of eye pigmentation) since the b-actin promoter is ubiquitously active and not Notch-restricted (Fig. S1). Incrossing TP1:Cas9-GFP, 4[U6:sg] F1s resulted in F2 adults with mosaic stripe mutations in patterns distinct from either F1 parent (Fig, 1c), suggesting TP1 drives early Cas9 expression in somatic cells and the transgene is active across generations. Collectively, these results demonstrate that CRISPR-Cas editing can be applied for recording signaling histories during zebrafish development.

**Figure 1.**
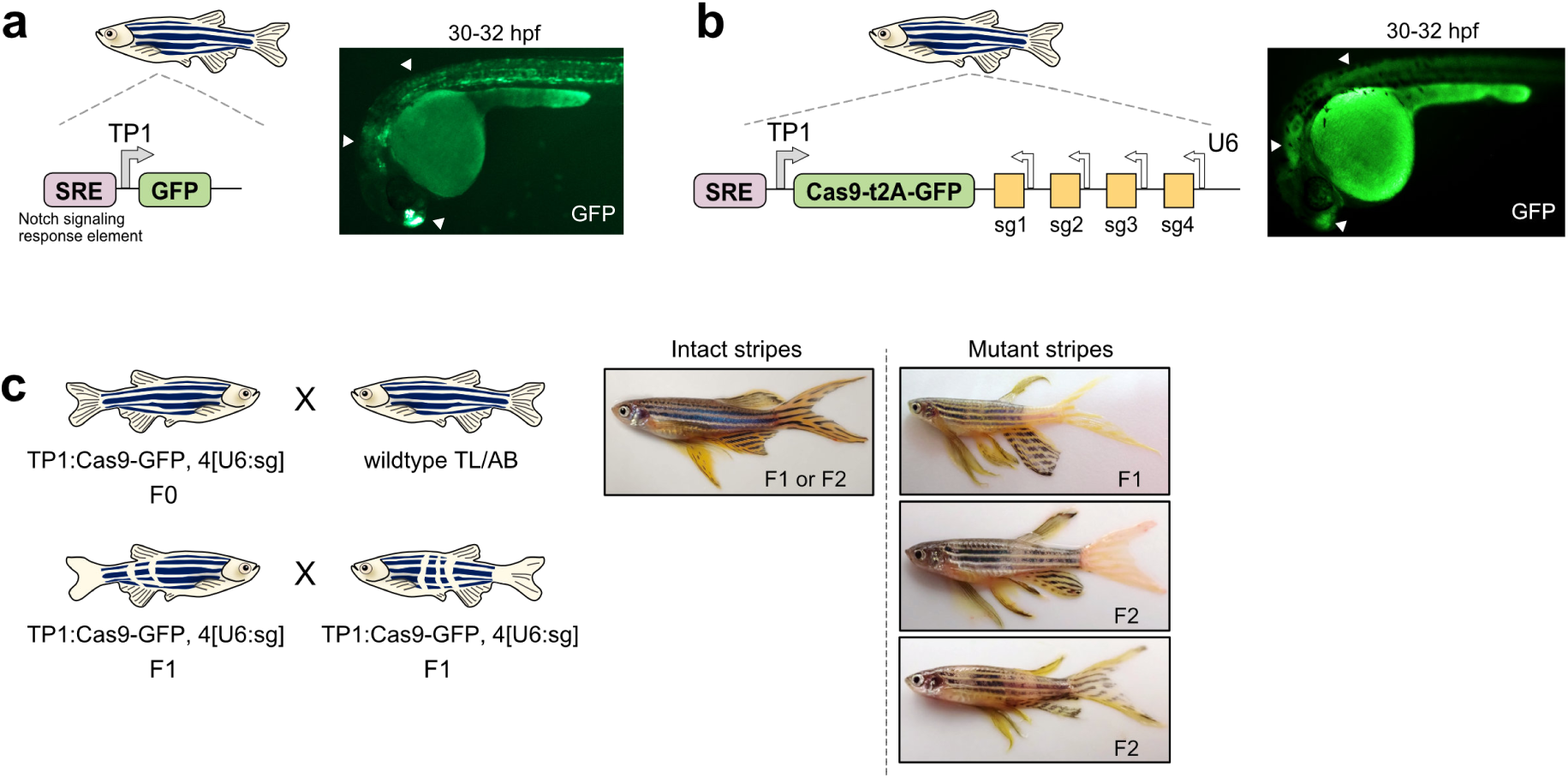
A Notch-dependent Cas9 zebrafish transgenic reporter. a) Left, schematic of TP1:GFP transgenic reporter. Right, fluorescence image of TP1:GFP expression at 30-32 hpf. Arrowheads highlight GFP expression in telencephalon, hindbrain and spinal cord. SRE, Notch signaling response element. b) Left, schematic of TP1:Cas9-GFP, 4[U6:sg] transgenic reporter. sg, sgRNA. sg4 targets *tyr* gene. Right, fluorescence image of TP1:Cas9-GFP, 4[U6:sg] expression at 30-32 hpf. Arrowheads highlight GFP expression in telencephalon, hindbrain and spinal cord. c) Left, schematics of TP1:Cas9-GFP, 4[U6:sg] outcross (top) and incross (bottom). Right, whole-body images of adult zebrafish with intact stripes and mutant stripes from F1 and F2 adults.

### SABER records Notch signaling in the developing zebrafish brain

To adapt CRISPR-Cas for Notch signal recording in the nervous system, we designed a new transgenic construct. Zebrafish *her4.3* (orthologous to mammalian *Hes5*) is highly expressed in neural progenitor cells and is a direct target of Notch signaling^29^. A 3.4 kb promoter fragment of *her4.3* containing five Notch co-factor binding sites (signaling response element) has been firmly established as a Notch activity reporter^30,31^. We generated transgenic zebrafish with a *her4.3* promoter driving dsRed and Cas9-GFP expression, and U6 promoters driving constitutive expression of four sgRNAs. This transgene, her4.3:switchCas9, represents a “Notch sensor”. To circumvent early silencing of transgenic Cas9 expression from constitutive promoters as discussed above, we flanked the dsRed sequence and a stop codon with loxP sites, such that Cas9-GFP will be “switched on” when both Notch signaling and Cre recombinase expression occur. This double inducible system enables recording of Notch signaling at timepoints of interest. Comparison of her4.3:GFP and her4.3:switchCas9 transgenic lines revealed similar expression patterns with fluorescence detected in the central nervous system (Fig. 2a, b). To confirm that her4.3:switchCas9 transgenics respond to Notch activity levels, we inhibited Notch signaling using the gamma secretase inhibitor LY411575^30,32,33^. We performed an early, short pulse inhibition (4 h treatment, 4-8 hpf) prior to the onset of her4.3 expression at ∼7.5-8 hp f^29^ as well as sustained inhibition (4-72 hpf). Both her4.3:GFP and her4.3:switchCas9 transgenics exhibited reduced fluorescence by 48 hpf from sustained inhibition with the greatest effect observed by 72 hpf (Fig. 2c, 2d). Short pulse inhibition resulted in decreased fluorescence in the her4.3:GFP transgenic but not in the her4.3:switchCas9 transgenic. We note that dsRed expression is not readily detectable in her4.3:switchCas9 embryos until 48 hpf, whereas her4.3:GFP embryos display strong GFP expression by 24 hpf. Differences in the fluorophores used and the larger construct size of her4.3:switchCas9 likely account for differences in relative intensities and the timing of expression observed with short pulse Notch inhibition. In contrast, performing Notch inhibition in Olactb:SpCas9-GFP, 4[U6:sg] transgenic embryos did not reduce GFP expression (Fig. S2a), demonstrating that her4.3:GFP and her4.3:switchCas9 transgenics specifically respond to Notch activity and confirming their functions as sensors.

**Figure 2.**
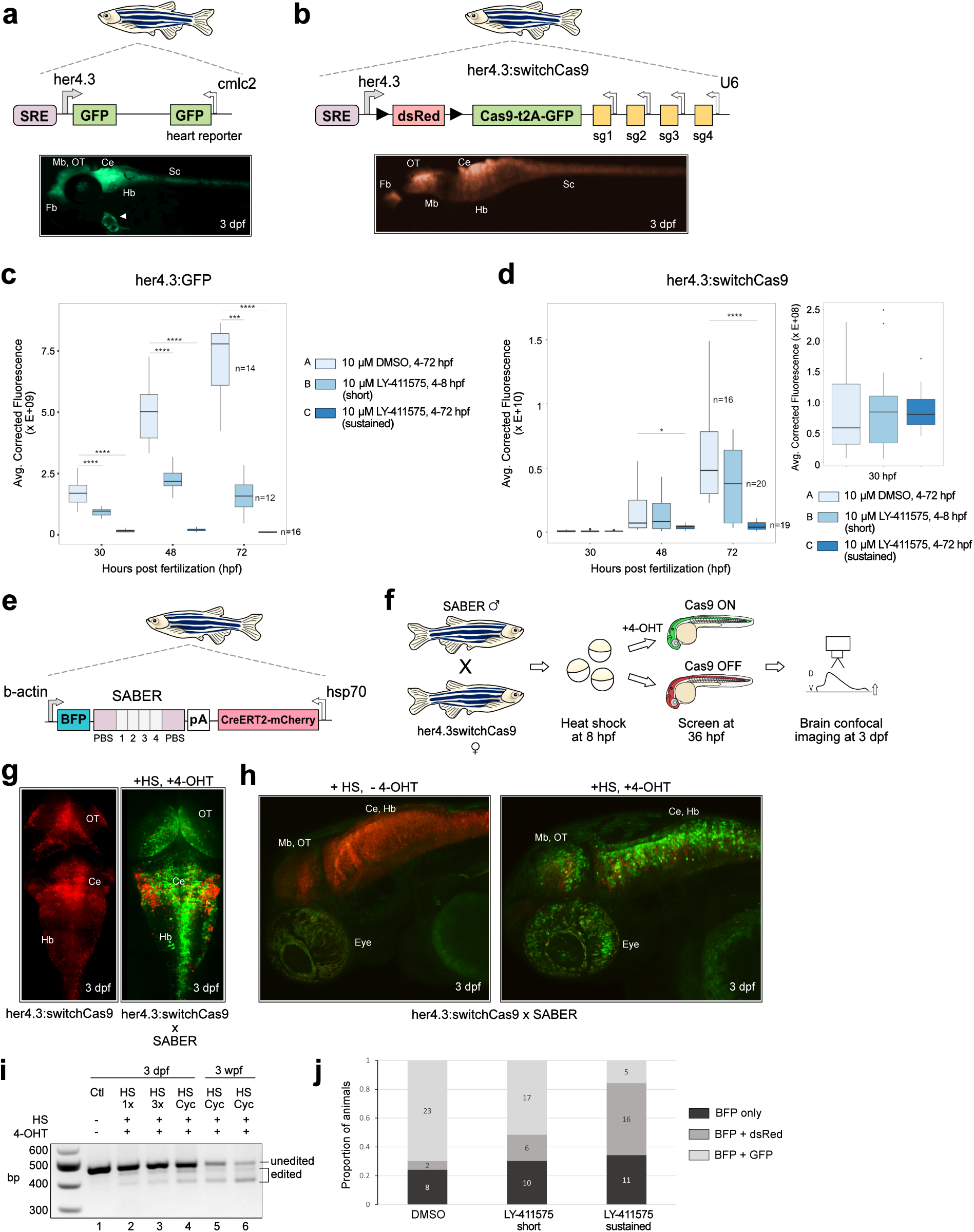
SABER, a novel CRISPR-Cas9 signal recorder. a) Top, schematic of her4.3:GFP, cmlc2:GFP transgenic reporter. Bottom, fluorescence image of her4.3:GFP expression at 3 dpf. Arrowhead, heart GFP expression driven by cmlc2. Fb, forebrain. OT, optic tectum. Mb, midbrain. Ce, cerebellum. Hb, hindbrain. Sc, spinal cord. b) Top, schematic of her4.3:switchCas9 transgenic reporter. sg, sgRNA. Bottom, fluorescence image of her4.3:switchCas9 expression at 3 dpf. c) GFP fluorescence intensity in her4.3:GFP embryos after Notch inhibition. Embryos were treated with DMSO control at 4-72 hpf, LY-411575 inhibitor at 4-8 hpf (short pulse), and LY-411575 inhibitor at 4-72 hpf (sustained). Images were taken at 30, 48 and 72 hpf. n, number of embryos. ****, p < 0.0001, Mann-Whitney U Test. d) dsRed fluorescence intensity in her4.3:switchCas9GFP embryos after Notch inhibition. Right panel, zoomed and rescaled graphs of dsRed intensity at 30 hpf. Identical treatment as embryos in (c). *, p < 0.05. ****, p < 0.0001, Mann-Whitney U Test. e) Schematic of SABER Notch transgenic recorder. BFP, blue fluorescent protein. PBS, primer binding sites in purple. Grey boxes 1-4, target sites matching sg1-sg4 sgRNAs on her4.3:switchCas9 transgene. Hsp70, heat shock promoter. f) Schematic of SABER Notch recording. SABER F1 male transgenic is crossed to her4.3:switchCas9 F1 female transgenic. F2 embryos are heat shocked at 37 °C for 30 min at 8 hpf to induce CreERT2 expression. A subset of embryos is treated with 4 hydroxytamoxifen (4-OHT) after heat shock until 3 dpf. Control embryos are not treated with 4-OHT. Confocal images are taken at 3 dpf. D, dorsal. V, ventral. g) Maximum projections showing dorsal brain confocal images of 3 dpf her4.3:switchCas9 transgenic embryos with dsRed expression (left) and her4.3:switchCas9 x SABER double transgenic embryos with GFP conversion after heat shock and 4-OHT addition (right). h) Maximum projections showing lateral brain and eye confocal images of her4.3:switchCas9 x SABER double transgenic embryos at 3 dpf after heat shock cycling without (left) and with (right) 4-OHT addition. i) Genomic DNA amplification of SABER to assess barcode editing efficiency. HS, heat shock. 1X, 1 heat shock pulse. 3X, 3 heat shock pulses. HS Cyc, heat shock cycling. wpf, weeks post fertilization. bp, base pairs. j) Proportion of 3 dpf animals from her4.3:switchCas9 x SABER cross with BFP, dsRed or GFP fluorescence after Notch inhibition. BFP only: barcode^+^ but dsRed/Cas9-GFP^-^. BFP + dsRed: barcode^+^ and dsRed/Cas9-GFP^+^ but no GFP detected. BFP + GFP: barcode^+^ and dsRed/Cas9-GFP^+^ and GFP detected. Numbers inside bars represent animal counts in each group.

To enable barcode recording, we designed a new transgenic construct with a b-actin promoter driving expression of blue fluorescent protein with a barcode array embedded in the 3’ UTR, and heat shock inducible expression of CreERT2 recombinase (Fig 2e). This transgene represents the “Notch recorder” henceforth referred to as SABER. The barcode is a concatemerized array of four target sites matching the four sgRNAs present on the her4.3:switchCas9 construct. To test Notch-dependent activation of Cas9, we crossed SABER and her4.3:switchCas9 transgenics, heat shocked the embryos at 8 hpf and added 4-hydroxytamoxifen (4-OHT) to a subset to trigger CreERT2-mediated loxP excision (Fig. 2f). Starting at 36 hpf, we screened for GFP expression as a proxy for Cas9 expression and performed confocal imaging at 3 days post fertilization (dpf). As expected, her4.3:switchCas9 embryos displayed red fluorescence (Fig. 2g). In contrast, SABER and her4.3:switchCas9 double transgenics had extensive conversion and GFP expression in the brain. We observed the highest GFP expression around ventricular zones of the telencephalon, optic tectum, cerebellum, and rostral hindbrain (Fig. 2g). We also observed GFP expression in the retina (Fig 2h). Raising double transgenic embryos to adults confirmed the presence of stripe mutations although the mutations were more subtle than those observed in TP1:Cas9-GFP, 4[U6:sg] adults (Fig. S2b). This is likely due to the more restricted expression pattern of her4.3 relative to TP1. To rule out effects of leakiness of the heat shock promoter, we compared brains from SABER x her4.3:switchCas9 crosses where all embryos were heat shocked but only a subset had 4-OHT added. In the absence of 4-OHT, we observed at most 1-2 cells that were GFP positive, confirming that Notch, CreERT2 and 4-OHT are required for activation of Cas9 (Fig. 2h).

Next, we determined if Notch-dependent Cas9 activation results in barcode editing. We extracted genomic DNA from the heads of 3 dpf single transgenic (BFP-barcode^+^) and double transgenic (Cas9-GFP^+^, BFP-barcode^+^) embryos and PCR amplified SABER barcodes. We observed no editing in embryos where Cas9 is not expressed whereas double transgenics treated with heat shock and 4-OHT displayed shifts in band sizes, indicating editing (Fig. 2i). Since Cre-mediated conversion efficiency can impact editing outcomes, we tested various heat shock conditions. We observed that a single 35 min heat shock pulse at 8 hpf (Fig 2f) resulted in a low level of editing (Fig. 2i, 1X HS). Increasing to 3 x 35 min pulses at 8, 24 and 32 hpf did not result in a noticeable difference (Fig. 2i, 3X HS). Next, we tested heat shock cycling (Fig. S2c). Embryos were subjected to a single 35 min heat shock pulse at 8 hpf and 4-OHT was added. At 24 hpf, once embryos were more robust, we transferred single embryos to PCR strip tubes, refreshed 4-OHT and performed 16 cycles of heat shock followed by recovery in a thermocycler. Overall, embryos received 17 heat shock pulses from 8-48 hpf. This resulted in enhanced barcode editing (Fig. 2i, HS Cyc). Notably, we observed that Cas9-GFP expression slowly faded 4-5 days after it was first activated, suggesting that signaling can be robustly recorded for a window of ∼96-120 hours after the initiation of heat shock and 4-OHT addition (data not shown). It is possible that very low levels of Cas9-GFP (undetectable by fluorescence) may persist for longer periods and enable continued signal recording, but this is likely to be rare. Furthermore, we extracted genomic DNA from the brains of 3 weeks post fertilization (3 wpf) double transgenic animals that were subjected to heat shock cycling as embryos and observed strong barcode editing (Fig. 2i). This demonstrates that signaling histories are permanently recorded and can be extracted long after the recording period has ended. Next, we tested the efficiency of switching on Cas9-GFP expression when Notch activity is inhibited. Performing a short pulse inhibition in embryos from SABER x her4.3:switchCas9 cross resulted in a slight reduction in the number of embryos with detectable GFP expression and no noticeable difference in barcode editing relative to DMSO controls (Fig. 2j, Fig S2d). In contrast, sustained Notch inhibition resulted in a dramatic reduction in the number of embryos expressing GFP and suppressed barcode editing (Fig. 2j, Fig S2d). Collectively, these results confirm that SABER can specifically and permanently record Notch activity in the brain in an inducible manner.

### SABER-seq recovers signaling histories and transcriptional states

Having established a Notch sensor (her4.3:switchCas9) and a Notch recorder (SABER), we implemented our goal of matching cell signaling histories in early development with the eventual cell fate identities adopted. We crossed SABER and her4.3:switchCas9 transgenics, heat shocked embryos and added 4-OHT at 8 hpf, performed heat shock cycling with fresh 4-OHT from 24-48 hpf, screened larvae for Cas9-GFP and BFP-barcode expression, and performed scRNA-seq of whole brains from 21-23 dpf juveniles using the 10X Chromium platform (Fig. 3a). This method, SABER-seq, enables simultaneous recovery of endogenous mRNAs and SABER barcode transcripts. We prepared SABER libraries via PCR amplification of barcode cDNAs and sequenced the libraries separate from corresponding transcriptome libraries. In our first iteration of SABER-seq, we recovered barcodes from 3,454 cells from two brains (b1, b2). This represents 6-8% recovery from all profiled cells per animal (version referred to as SABER_v0). To improve recovery, we optimized SABER-seq library preparation steps (see Methods) and recovered 3,156 cells from a third brain (b4), which represents 22% recovery. (version referred to as SABER_v1). In summary, we obtained 6,610 barcodes from single cells of which 5,351 barcodes (81%) were mappable to the unedited reference sequence (Table S1). The remaining barcodes were too short (less than 50 bp) for unambiguous mapping. Large deletions resulting in very short amplicons have been previously described as a limitation of Cas9 editing^34^. To analyze the spectrum of mutations, we only considered the mappable cohort of barcodes for downstream analyses. As expected, barcode editing was highest around the sgRNA target sites but extended upstream and downstream of the targets, consistent with larger deletions (Fig. 3b). Furthermore, 83.4% of barcodes had edits in at least one target site, confirming effective Cas9 expression and activity in Notch induced cells. Deletions were the predominant mutations and could be categorized as intra-site - edits within a target - and inter-site, edits that span two or more targets or extend outside the sgRNA loci (Fig. 3c, 3d). The frequency of each type of deletion varied among the four target sites with deletions spanning target sites 1 to 4 being the most frequent (Fig. 3d).

**Figure 3.**
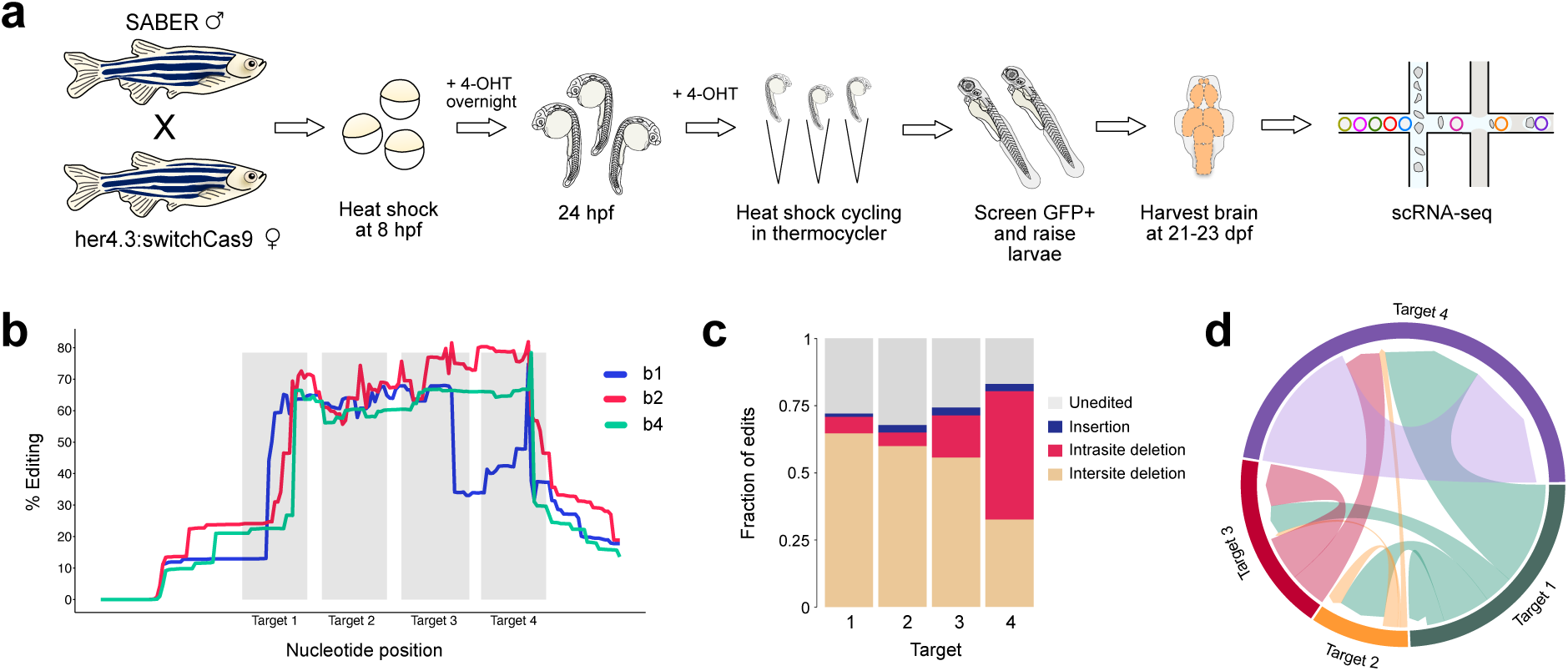
SABER-seq recovers Notch-edited barcodes from single cells. a) Schematic of SABER-seq experimental protocol. b) Percent editing across each nucleotide in SABER-seq barcodes. The positions of the four CRISPR target sites are indicated in grey. Data from three individual brains is shown. Only deletion mutations are shown. c) Edit type at each target site. d) Chord diagram of the nature and frequency of deletions. Each colored sector represents a target site. Links between target sites indicate inter-site deletions; self-links indicate intra-site deletions. Link widths are proportional to edit frequencies.

As discussed above, no current methodology enables high resolution, high throughput mapping of cellular signaling to distinct molecular cell types in complex tissues during development. To superimpose Notch signaling histories captured via SABER-seq onto finely resolved brain cell types and cell states, we first established a juvenile zebrafish brain cell catalog. In previous work^7^, we used inDrops to describe brain cell taxonomy from similar aged animals (23-25 dpf) but to avoid technical artifacts of mapping SABER-seq data obtained via the 10X Chromium platform onto pre-existing inDrops data, we collected new samples. We profiled brain regions (telencephalon, diencephalon/midbrain, and hindbrain tissues) and whole brains (including brains b1, b2, b4 from SABER-seq experiments) from 21-23 dpf animals using 10X Chromium. We sequenced 156,353 cells and retained 148,853 cells for downstream analysis after quality filtering. Using a combination of initial coarse-grained computational clustering and subsequent rounds of iterative clustering, we classified 137 neuronal, non-neuronal and progenitor cell subtypes and cell states (Fig. 4, Fig. S3, Table S2). This represents more annotations than identified in our previous dataset^7^ (137 vs 105), which is expected since we sequenced greater than twice the number of cells. For example, we identified 10 granule cell subclusters, 12 pallium subclusters, 6 subpallium subclusters, 9 hypothalamus subclusters, 9 dorsal habenula subclusters, 6 ventral habenula subclusters, and 7 radial glia subclusters, among many others. Although we used classical markers from literature to assign subtypes to anatomical regions as described previously^7,35^, a few cell types could not be confidently mapped to discrete regions and were labeled by their enriched cluster marker only (e.g. Neuron_mef2cb^+^). There were also two clusters whose identities we could not fully resolve (Unk_1, Unk_2). We observed considerable overlap between markers identified in our study and those used to classify and in vivo validate subtypes described in recent scRNA-seq studies where corresponding brain subregions were dissected for profiling^36–38^. For example, a scRNA-seq study^37^ of pallium (*eomesa*, *bhlhe22*, *zbtbt18*, *emx3* expressing) and subpallium (*dlx2a*, *dlx5a*, *dlx6a* expressing) tissues identified cells expressing *c1ql3b.1* (Pal_2), *lhx9* (Pal_3), *pvalb7* (Pal_11), *otpa* (SubPal_1), *lhx6* and *sst1.1* (SubPal_4), and *six3b* (SubPal_6), among others (Fig. S3c, S3e). A scRNA-seq study^38^ of *gng8^+^* habenula cells validated cells expressing *wnt7aa* (cluster DHab_4), *adcyap1a* (DHab_5), *foxa1* (DHab_6), and *cbln2b* (DHab_8) among others (Fig. S3f). In addition, a scRNA-seq study^36^ of the preoptic area and hypothalamus identified cells expressing *trh* (POA_2), *sp8a* (Hyp_5), *prdx1* (Hyp_6), and *gsx1* (Hyp_7) among others (Fig. 4c). Finally, a scRNA-seq study^10^ of radial glia (*fabp7a*, *glula*, *cx43* expressing) identified cells expressing high levels of *her4.3* (RG_2) and *robo4* (RG_3) (Fig. 4d). Collectively, we observe that our whole brain dataset contains fewer subtypes than those obtained from enriched profiling of individual brain subregions. This is expected as we have less than 1X coverage of 21-23 dpf brain cells (estimated to have >1 million cells). Nevertheless, we identified multiple rare populations including pineal cells (56 cells representing 0.04% of dataset) and olfactory bulb cells (80 cells representing 0.05% of dataset) (Fig. S3h, Table S2).

**Figure 4.**
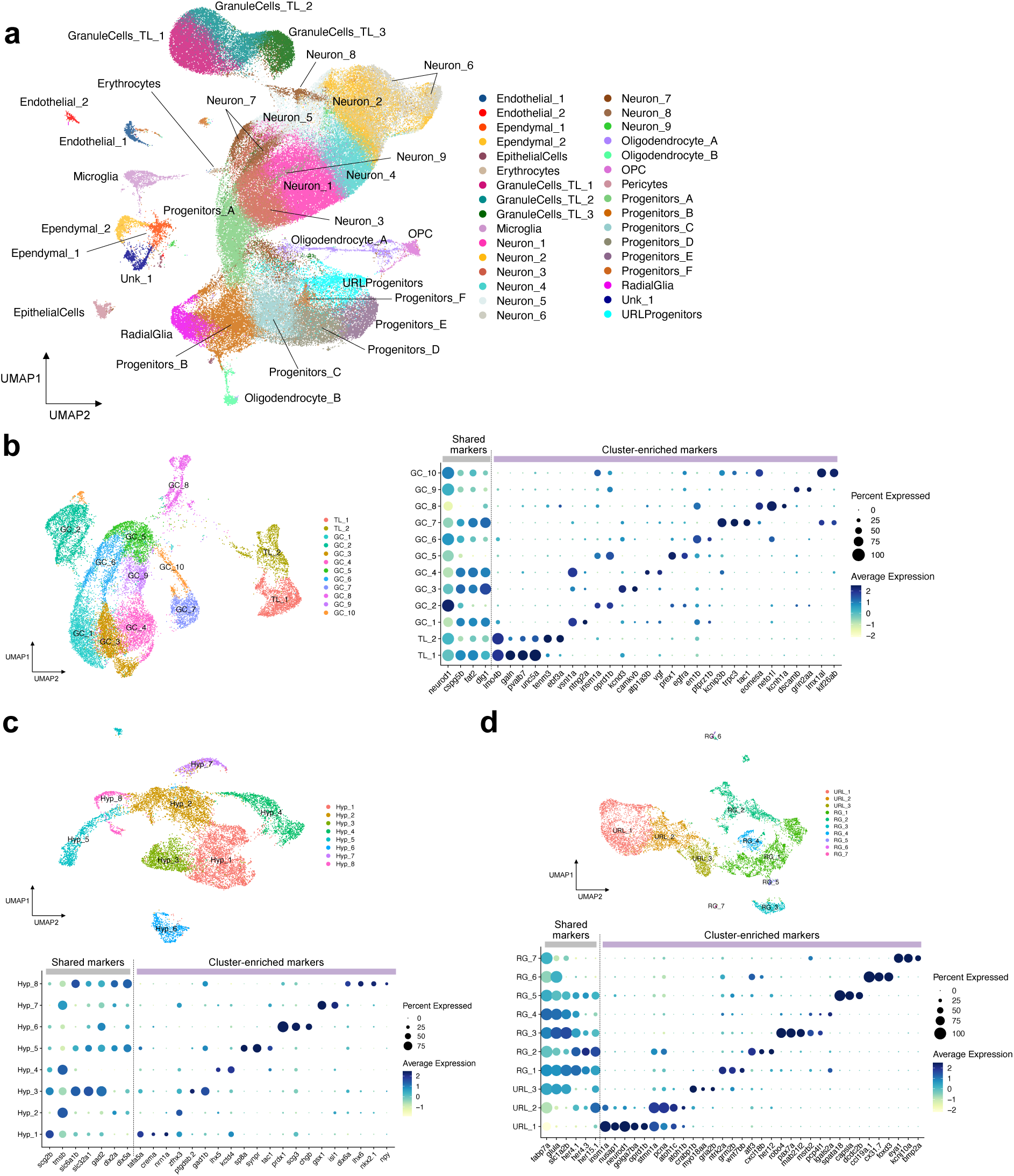
An expanded juvenile zebrafish brain atlas. a) UMAP embedding of 148,853 cells showing 32 clusters resulting from coarse-grained clustering. UMAP embedding and dot plots of marker gene expression from subclustering of b) granule cells and torus longitudinalis cells, c) hypothalamus cells, d) radial glia and upper rhombic lip progenitor cells. Note, hypothalamus had another subcluster identified not shown in (d), see Fig. S3h.

### Signal tracing reveals that Notch stimulated progenitors give rise to the majority of zebrafish brain cell types

Next, we matched SABER-seq barcodes with our annotated brain cell dataset. We obtained 277 uniquely edited SABER barcodes of varying frequencies. The most abundant barcode comprised 23% of profiled cells and was found in brain samples from all three animals, suggesting barcode homoplasy (i.e. identical barcodes arising in parallel independently) occurs with SABER-seq. Barcode homoplasy has been previously reported with CRISPR-Cas lineage recorders that rely on in vivo expression of Cas9 in zebrafish^7^. Although barcode homoplasy confounds clonal tracing (i.e. identifying common ancestors), it is not a limitation when barcodes are used to investigate signaling histories. Our “signal tracing” method is designed to determine the fraction of a profiled sample that is derived from progenitors where detectable signaling occurred (Fig. 5a). SABER-seq barcodes, including “unedited” barcodes, overlapped 127/137 cell annotations in our dataset (Fig. 5b, Table S3). The missing cell types were rare populations comprising less than 0.12% of the dataset except for two forebrain subclusters (Di_4 and Tel) that represented 0.2% of the data. Thus, barcodes were likely not detected in those cells due to lack of sampling depth. Importantly, edited SABER barcodes were detected in 126/127 subclusters from which barcodes were recovered. This suggests that detectable Notch signaling, as measured by Cas9 editing, occurred in nearly all recovered cell types and states and that Notch signaling is a component of the developmental histories of most brain cell types. We asked whether cells of a given cell type are detected more or less often in the SABER-seq dataset relative to the rest of the dataset. We identified 21 subclusters where cells were detected more frequently and 22 subclusters where cells were detected at a lower frequency (Table S3). These results suggest some bias in the expression and/or amplification of the Notch recorder transgene, which is driven by a beta-actin promoter, among the different cell types.

**Figure 5.**
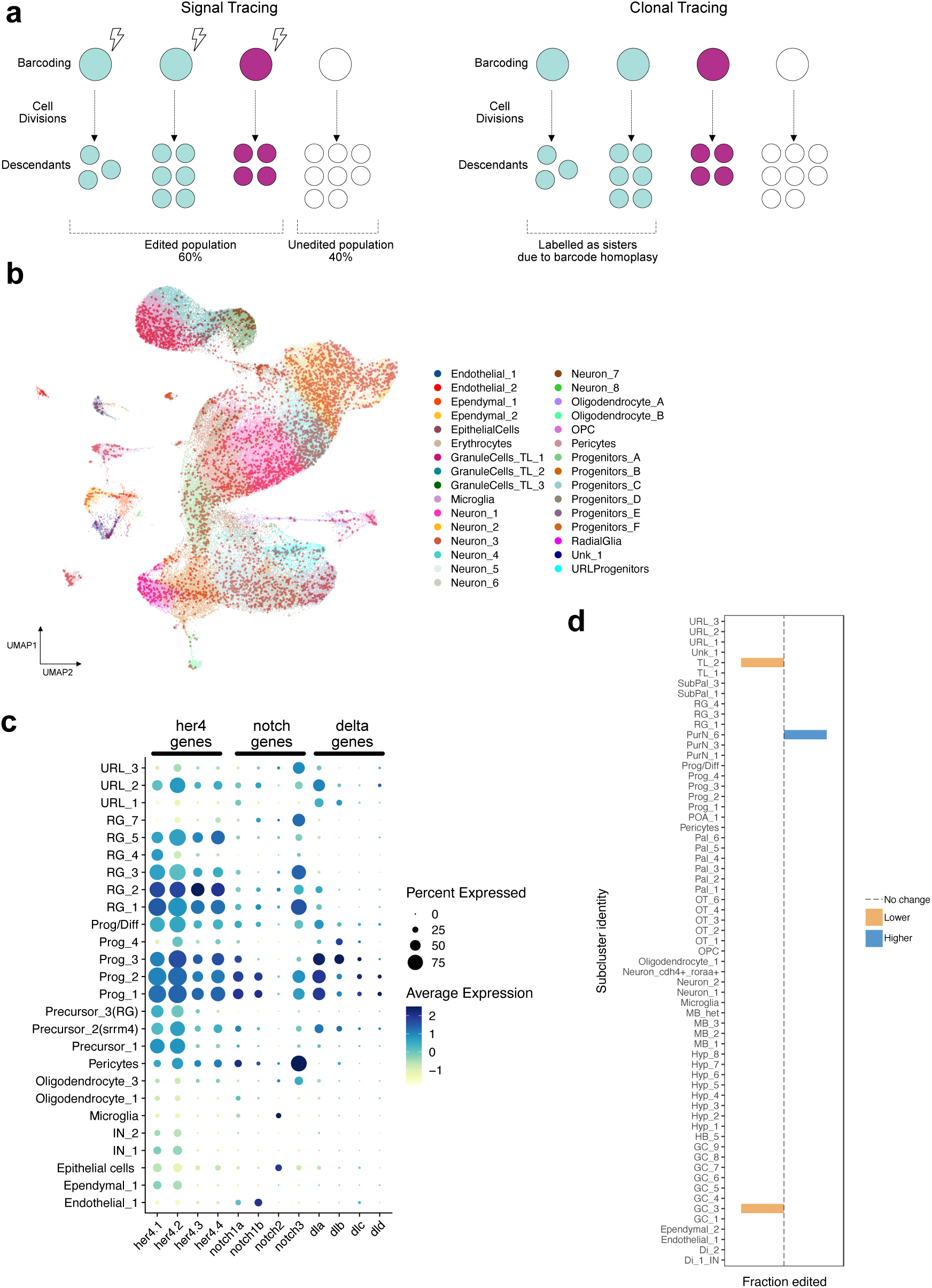
SABER-seq barcodes matched to cell type atlas. a) Schematic of difference between signal tracing and clonal tracing. b) UMAP embedding of 148,853 cells from Fig. 4a with cells from which SABER barcodes were recovered highlighted in red. c) Gene expression dot plot of her4 variants, notch genes and delta genes in the indicated subclusters. d) Comparison of fraction of cells with edited versus unedited SABER barcodes across subclusters where at least 20 cells had associated SABER barcodes. Dotted line indicates no significant change in proportion of edited vs unedited barcodes relative to all other clusters. Blue bar indicates higher proportion of edited vs unedited barcodes than expected (p < 0.05, binomial test). Orange bar indicates lower proportion of edited vs unedited barcodes than expected (p < 0.05, binomial test).

We investigated whether Notch recording occurred in the expected cell types in our dataset. Since Cas9 expression was initiated in cells expressing her4.3:switchCas9 upon Notch activation, edited SABER barcodes should be detected in the progeny of Notch stimulated-her4.3^+^ ancestor cells, effectively acting as a permanent tracer of descendants. The zebrafish *her4* gene cluster is organized as tandem duplicate repeats (her4.1-her4.5) on chromosome 23 with nearly identical transcripts, except for some polymorphisms in the 3’ UTR. Additionally, all variants are translated into identical peptides. We inspected the expression of endogenous her4 variants in our dataset. We observed highest expression of her4 variants including *her4.3* in several clusters of radial glia, which have neural stem cell properties, and other neural progenitor and precursor cell clusters in our dataset at 21-23 dpf (Fig. 5c, Fig. S4). This is consistent with the constitutive neurogenic capacity of these cells at all stages from embryo to adult^39^. Furthermore, these cell types give rise to nearly all neuronal subtypes in the zebrafish brain^39^. Accordingly, we detect edited SABER barcodes in neural progenitors, immature neurons and mature neurons (Fig. 5b). Interestingly, we also recovered edited SABER barcodes from several classes of differentiated non-neuronal cells including epithelial cells, ependymal cells, oligodendrocytes and pericytes. Apart from pericytes, *her4.3* transcripts are barely detectable in these cell types at 21-23 dpf (Fig. 5c) although *her4.1* and *her4.2* transcripts are present. However, it is likely that *her4.3* was transiently expressed earlier during fate specification of these cell types before being shut off during or after differentiation, similar to the patterns observed in the transition from neural progenitors to neurons. Consistent with this, oligodendrocytes and ependymal cells have been identified as *her4.3*-derived progeny from lineage tracing experiments using scRNA-seq^40,41^. Thus, although *her4.3* expression at the time of cell isolation at 21-23 dpf does not fully resemble her4.3:switchCas9 transgene expression in all ancestral cells when the inherited SABER barcodes were generated, our data and past literature^31,40,41^ strongly suggest that it can be used as a proxy to identify Notch-active founder cell types and trace their descendants at later stages. Notably, Notch signaling has been shown to be important for cell fate specification of the aforementioned non-neuronal cell types in development and regeneration^42–46^. Regulation of *her4.3* expression is orchestrated via *notch1a*, *notch1b* and *notch3* receptor genes^31^. Indeed, expression of at least one of these notch genes was detected in cell types expressing *her4.3*, with *notch3* showing the highest levels and percent expression in our dataset (Fig. 5c). Among Notch ligands, Delta has been implicated in Notch signaling in the zebrafish brain^47^. In our dataset, *dla* had the strongest expression and overlapped with notch genes in the cell types where *her4.3* was highly expressed. Collectively, our results indicate that widespread Notch and *her4.3* activity in multiple cell types in the brain can be effectively captured by our technology.

Finally, we asked whether edited barcodes were detected more or less frequently than unedited barcodes in any of the identified subclusters, suggesting that Notch signaling in certain progenitor or precursor states may have differential impact on the terminal cell fates adopted (Fig. 5d). We only considered subclusters where at least 20 cells had associated barcodes to avoid low sampling bias. We identified two subclusters, GC_3 and TL_2, corresponding to a granule cell subtype and a torus longitudinalis subtype with lower proportions of edited barcodes than expected. In contrast, one subcluster, PurN_6, corresponding to a Purkinje cell subtype had a higher proportion of edited barcodes than expected (100% editing). These results suggest that development of GC_3 and TL_2 cells may be less sensitive or dependent on Notch signaling during early stages of development (< 5 dpf) when SABER recording occurred. On the other hand, Notch signaling may be fundamental for development of PurN_6 cells during the same temporal window. Importantly, these results demonstrate the power and sensitivity of SABER-seq for converting transient signaling events to permanent traceable marks and for identification of cells whose fates are determined downstream of these signaling activities.

## DISCUSSION

In the past few years, DNA molecular recorders have rapidly become one of the leading methods for massively parallel recording of various cellular paradigms owing to their scalability and compatibility with genomics, especially scRNA-seq. Nevertheless, most applications of such recorders for studying in vivo development have been limited to lineage tracing. Here, we demonstrate the potential of using a molecular recorder for signal tracing during zebrafish brain development. The power of SABER-seq rests on engineering transient Notch cellular signals to trigger permanent CRISPR/Cas-mediated edits in a genomic transgenic array, which are then propagated during cell division, enabling long term storage of cellular behavior. The recording is inducible and can be initiated at developmental windows of interest using 4-OHT and heat shock treatments. Additionally, SABER-seq enables simultaneous recovery of cellular transcriptomes to identify individual cells and cell types that develop post Notch activation. We also demonstrate the modular nature of SABER-seq using two different Notch sensitive promoters, TP1 and her4.3 in zebrafish, which were conveniently swapped via Gateway cloning. Thus, we expect that SABER-seq can be adapted for measuring additional signaling pathways by using a promoter with the appropriate upstream signaling response element.

We were surprised to observe stripe mutations in adult fish resulting from the cross of SABER and her4.3:switchCas9 (Fig. S2). The mutations indicate that her4.3:switchCas9 was expressed in ancestors of melanophores. Although expression of the her4.3 promoter fragment is well documented in neural tissues, its expression patterns in other regions are not well characterized. Nevertheless, studies utilizing the her4.3 transgene, her4.3:GFP, in non-neuronal contexts have shown that it is active in neural crest cells as well as in blastemas during fin regeneration^30,48^, warranting a more thorough exploration of her4.3 transgene expression. Thus, scRNA-seq of SABER barcodes in other tissues may reveal previously unrecognized sites of transient *her4.3* and Notch activities. Our results also suggest that some terminal cell fates in the brain may be more or less closely linked with Notch signaling than others. It remains to be determined if these effects are directly regulated by Notch, however our findings demonstrate how SABER-seq can be used to begin to explore the relationship between signaling and cell fate decisions.

SABER-seq provides several advantages over previous iterations of signaling reporters. First, since it is a sequencing-based method, it is scalable to millions of cells in a cost-effective manner. We describe its application with 10X Chromium, but we anticipate it can be combined with other lower cost scRNA-seq platforms such as sci-seq^49^ by simply adapting the SABER library amplification protocol. Second, the permanence of the barcodes precludes the need for real time observation of signaling activity or being constrained to short term observation, issues which plague fluorescent signal recorders where fluorescence is detectable for limited number of cell divisions. Third, it provides a detailed measurement of cellular gene expression states while fluorescent reporters lack this ability. These features make SABER-seq an attractive alternative to traditional signaling reporters, however we foresee several optimizations. First, barcode recovery is currently limited to less than 25% of profiled cells. Differences in barcode expression between cell types and short amplicons resulting from large CRISPR/Cas deletions that often remove primer binding sites introduce bottlenecks in barcode library generation^7^. To circumvent the latter, fusion proteins of Cas9 and DNA nucleotidylexotransferase (DNTT, also known as TdT), a template-independent polymerase, can be used to favor sequential insertions and enhance barcode recovery^22^. Second, SABER-seq is currently a binary system where barcodes are either edited (signal detected) or unedited (no signal or insufficient signal). However, differences in signaling strength based on levels, duration, frequency or oscillations of Notch signaling cannot be reliably converted into meaningful differences in barcode editing frequencies. Finally, scRNA-seq results in tissue destruction and loss of spatial context. Although, it is possible to perform in silico mapping of cells to their most likely location using spatial markers, it is still prediction based. Furthermore, although clustering is powerful when combining many single cells together, matching SABER-seq barcodes with transcriptional identities is done at the level of individual cells. Thus, if an individual cell is erroneously grouped with other similar but not identical cells in a given cluster, it will be misclassified. To mitigate these drawbacks, we anticipate merging SABER-seq with spatial transcriptomics^50^ in the future to preserve tissue architecture and niche information, and aid cell type classification.

SABER-seq measures a different paradigm than lineage molecular recorders. The latter is designed to determine cell division patterns and relatedness between cells. As a result, high barcode diversity and low barcode homoplasy are required for unambiguous mapping of clonal relationships. In contrast, SABER-seq is engineered to determine the fraction of a population arising from signal-activated ancestral progenitors. For signals like Notch that are active in many cells and for long temporal windows, barcode editing is expected in a higher fraction of the profiled cells. For signals with more restricted spatiotemporal patterns, a lower fraction of profiled cells will display barcode editing. Thus, although descendants of signal-activated ancestors are marked, the relationship between marked cells is not analyzed by signal tracing. By varying the timing of signal recording, multiple signal tracing snapshots can be stitched together to track variations in signaling patterns throughout development via changes in the barcoded cell populations. To merge SABER-seq with lineage tracing, barcode complexity will need to be enhanced by shifting to Cas9-DNTT fusions for editing as well as by increasing the target sgRNA loci from four as in this study to at least 8-10 sites as used in other lineage tracing technologies^7,22,35,51^.

We propose some exciting applications of SABER-seq in its current format. First, it can be used to investigate the role of signaling on fate specification by perturbing signaling using chemical inhibitors or genetic knockdowns or knockouts of signaling components and assessing their effects on the composition of cell populations via signal tracing maps. Such “signalomes” can provide a relatively rapid, high throughput, and unbiased screening method to identify signaling effectors that impact discrete cell fate decisions for downstream functional validations. Second, it can be used to study changes in signaling activity accompanying cell fate remodeling during regeneration of limbs or organs as well as during oncogenesis and metastases.

In summary, SABER-seq lays the foundation for massively parallel signal recording with single cell resolution. Our proof-of-principle study describing its application for sensing and storing Notch signaling inputs during brain organogenesis establishes another use of CRISPR/Cas molecular recorders in development. We anticipate further design adaptions of this technology for multiplexed recording of two or more different signaling pathways or multiplexed temporal recording of the same pathway using biochemically diverse orthologous Cas proteins.

## ACKNOWLEDGEMENTS

We thank Hemagowri Veeravenkatasubramanian for discussion, advice and comments on the manuscript. We thank Laure Bally-Cuif for sharing various her4.3 plasmid constructs, Diego Jimenez for suggesting the SABER acronym, the UPenn zebrafish facility staff for assistance with animal husbandry, and Michael Granato and the UPenn Cell and Developmental Biology microscopy core for use of their confocal microscopes. This work was supported by DGE-2236662 NSF GRFP to J.E.G. and R00HD098298 and DP2NS131787 to B.R. Any opinions, findings, and conclusions or recommendations expressed in this material are those of the author(s) and do not necessarily reflect the views of the National Science Foundation.

## AUTHOR CONTRIBUTIONS

B.R. conceived and designed the study and wrote the manuscript. A.S. R.P.P. and J.E.G. edited the manuscript. A.S., A.M., and H.D. cloned constructs and/or generated transgenic lines. A.S. performed Notch inhibition, confocal imaging and Notch signal recording experiments. A.M., J.E.G. and B.R. performed scRNA-seq experiments. B.R. performed SABER-seq experiments. R.P.P. processed the SABER-seq pipeline. R.P.P., J.E.G. and B.R. performed bioinformatic data processing.

## COMPETING FINANCIAL INTERESTS

The authors declare no competing interests.

## METHODS

### Zebrafish husbandry

This work was performed under protocol numbers 807110 and 807259, which were approved by the University of Pennsylvania’s Office of Animal Welfare of Institutional Animal Care and Use Committee (IACUC). All zebrafish work in this study follows the University of Pennsylvania Institutional Animal Care and Use Committee regulations.

### Constructs for transgenesis

The plasmid pTol2-her4.3-eGFP-cmlc2-GFP was a gift from Laure Bally-Cuif. The plasmid pTol2-TP1bglob-eGFP (TP1:GFP in this study) was obtained from Addgene (plasmid 73586). The plasmid pTol2-TP1-SpCas9-t2A-GFP, 4xU6:sgRNA (TP1:Cas9-GFP, 4[U6:sg]) and pTol2-her4.3:loxP-dsRed-loxP-SpCas9-t2A-GFP, 4xU6:sgRNA (referred to as her4.3:switchCas9) were constructed as follows. Individual sgRNAs targeting sites 1-4 of the SABER array were cloned into 4 separate U6x:sgRNA plasmids (Addgene plasmids 64245-64248) as described previously^52^. The U6x:sgRNAs were assembled into a contiguous sequence in the pGGDestTol2LC-4sgRNA vector (Addgene plasmid 64242) by Golden Gate ligation. The resulting 4xU6:sgRNA sequence was PCR amplified and ligated into the backbone of pDestTol2pA2-U6:gRNA (Addgene plasmid 63157) after the vector was first digested with ClaI and KpnI to generate the pDestTol2pA2-4xU6:sgRNA plasmid. The 5’ entry vector p5E her4.3-loxP-dsRed-SV40pA-loxP was generated by first PCR amplifying the loxP-dsRed-SV40pA-loxP sequence from pTol2Olactb:loxP-dsR2-loxP-EGFP vector (gift from Atsushi Kawakami). The amplified sequence was then ligated into the backbone of plasmid p5E-her4.3 (gift from Laure Bally-Cuif) after the vector was digested with SalI and NotI. The final constructs were generated via multisite Gateway with p5E-TP1 (Addgene plasmid 73585) or p5E her4.3-loxP-dsRed-SV40pA-loxP together with pME-Cas9-t2A-GFP (Addgene plasmid 63155), p3E-polyA (Tol2 kit) and pDestTol2pA2-4xU6:sgRNA.

The SABER barcode transgenic vector pTol2-Olactb-BFP-SABER-SV40-hsp70-mCherry-CreERT2-SV40 was constructed as follows. The SABER barcode sequence was ordered as a gene block from IDT and was cloned into a multiple cloning site (MCS) in the 3’ UTR of a BFP coding sequence in the base vector pTol2-Olactb-BFP-MCS-SV40-hsp70-mCherry-CreERT2-SV40. Plasmids will be available from Addgene.

### Generation of transgenic zebrafish

All transgenic lines were generated by injecting one-cell embryos with Tol2 mRNA and corresponding plasmids, identifying founders and raising F1 adults as described previously^53^.

### Notch recording

Male SABER adults were crossed to female her4.3:switchCas9 adults. For single heat shock treatment, 8 hpf embryos were heat shocked for 35 minutes at 37 °C. Embryos were transferred to a petri dish of fresh media containing 10 μM 4-OHT (4-hydroxytamoxifen) and incubated at 28.5 °C until screening at 2-3 dpf. For triple heat shock treatment, embryos were first heat shocked at 8 hpf and 4-OHT added as described above. Next, embryo heat shock was repeated at 24 hpf and fresh 4-OHT was added. Finally, embryo heat shock was repeated at 48 hpf and fresh 4-OHT was added. Larvae were screened at 3 dpf. For heat shock cycling, embryos were heat shocked at 8 hpf for 35 minutes at 37 °C and transferred to a petri dish of fresh media containing 10 μM 4-OHT for overnight incubation at 28.5 °C. At 24 hpf, embryos were screened for whole body BFP expression to identify the SABER transgene. Positive embryos were transferred to individual PCR tubes containing 100 μl 10 μM 4-OHT. The embryos were heat shocked in a thermocycler for a total of 16 cycles of 25 minutes at 37 °C and 45 minutes at 28.5 °C. After the first four cycles, 4-OHT was refreshed, and the fish were returned to the thermocycler to complete treatment. Following the last cycle (∼48 hpf), larvae were transferred to petri dishes containing 10 μM 4-OHT and incubated at 28.5 °C for four hours after which larvae were screened for GFP and dsRed.

### Notch inhibition

her4.3:GFP, her4.3:switchCas9 and Olactb:SpCas9-GFP, 4[U6:sg] embryos were treated with 10 µM LY-411575 or dimethyl sulfoxide (DMSO) from 4 to 8 hpf or 4 to 72 hpf and imaged at 30, 48, and 72 hpf. After imaging, larvae were transferred to individual wells of a 24-well plate for continuous identification. Notch inhibition in SABER x her4.3:switchCas9 fish was performed similarly with the following modifications. At 8hpf, embryos were heat shocked for 35 minutes at 37 °C and transferred to a petri dish of fresh media containing 10 μM 4-OHT and either 10µM LY-411575 or DMSO and incubated overnight at 28.5 °C. At 24 hpf, embryos were screened for whole body BFP expression to identify the SABER transgene. Positive embryos were transferred to individual PCR tubes containing 100 μl 10 μM 4-OHT and either 10 µM LY-411575 or DMSO. The embryos were heat shocked in a thermocycler for a total of 16 cycles of 25 minutes at 37 °C and 45 minutes at 28.5 °C. After the first four cycles, the drugs and media were refreshed, and the fish were returned to the thermocycler to continue cycling. Following the last cycle (∼48 hpf), the larvae were transferred to petri dishes containing 10 μM 4-OHT and either 10 µM LY-411575 or DMSO and incubated at 28.5 °C for four hours after which larvae were screened for GFP and dsRed.

### Imaging

Fluorescence imaging was performed using a Leica M165 FC. Fluorescence intensity was analyzed in FIJI (Image J) 2.9.0. Corrected Fluorescence (CF) of the larval brain was measured by subtracting the product of the selection area and the mean grey value of the background from the integrated density of the selection area in each image. The Average CF (ACF) was calculated for each time point and treatment group. Confocal imaging was performed using a 40X water immersion lens on an Olympus Spinning disk confocal microscope and 3i Slidebook Software or a 20X lens on a Zeiss LSM 880 laser scanning confocal microscope. Images of adult fish were taken with a digital camera. To analyze differences in fluorescence intensities in Notch-inhibited groups a Mann-Whitney U Test was used.

### Whole brain scRNA-seq

Wild type and SABER x her4.3:switchCas9 zebrafish brains were harvested and dissociated at 21-23 dpf as described previously^53^ with the following modifications. All microcentrifuge tubes were treated with 1% BSA/DPBS overnight to prevent cell loss due to adhesion. Brains were dissociated with 1 mL of 20 units/ml papain in EBSS media and incubated at 37 °C for 20 min. Cells were resuspended with ∼150 μl ice cold 1%BSA/DPBS solution. Samples were run on the 10X Chromium scRNA-seq platform according to the manufacturer’s instructions (Single Cell 3’ v3 kit). Libraries were processed according to the manufacturer’s instructions. Transcriptome libraries were sequenced using HiSeq S4 300 cycle kits (Novogene) and Novaseq SP 100 and 200 cycle kits (MedGenome). Note, single brains were used for all experiments with whole brain samples. In contrast, 8-10 brains were pooled for experiments where forebrain, midbrain and hindbrain regions were dissected and profiled separately.

### SABER-seq

Notch recording was performed as described above. Animals with GFP expression were raised to 21-23 dpf in a fish facility. Individual brains were dissected and dissociated as described above. To generate SABER-seq libraries, samples post cDNA amplification and prior to fragmentation were split into two parts. One part was processed for transcriptome libraries as instructed by the manufacturer. The other part was processed for SABER-seq libraries as follows. To enrich for SABER barcodes, 5 µl of the whole transcriptome cDNA was PCR amplified with 10X_p140v1UP (GTGACTGGAGTTCAGACGTGTGCTCTTCCGATCT CGTCGTAGATCTTCATCGTGATG) and 10XPCR1F (CTACACGACGCTCTTCCGATCT) primers and Q5 polymerase. The reaction conditions were: 98 °C, 30 s; [98 °C, 10 s; 68 °C, 25 s; 72 °C, 15 s] x 14 cycles; 72 °C, 2 min. The reaction was cleaned up with 0.8X SPRISelect beads and eluted in 20 µl elution buffer (EB) buffer. A second PCR was carried out using 3 µl of PCR1 product and 10XPCR1F and 10X_p140v2Sa (GTGACTGGAGTTCAGACGTGTGCTCTTCCGATCT GAAACTACATGGGGTcaGCTC) primers. Reaction conditions were same as PCR1 except only 13 cycles were completed. At this point, 2 different clean up conditions were used: iteration 1 (SABER-seq_v0) and iteration 2 (SABER-seq_v1). For iteration 1, the reaction was cleaned up with 0.8X SPRISelect beads and eluted in 20 µl elution buffer (EB) buffer similar to PCR1. For iteration 2, an optimization was trialed, and the reaction was cleaned up with right side size selection using SPRIselect beads (0.6X beads used initially and supernatant was rebound with 1.5X beads). The product was eluted in 20 µl EB. Finally, adapters and sample indices were incorporated in a third PCR reaction using 5-7 µl of a 1:10 dilution of PCR2 product. Reactions conditions were same as PCR1 except 9-10 cycles were completed. Primers used: 10XP5Dual_Hx (AATGATACGGCGACCACCGAGATCTACAC-xxxxxxx xxx-ACACTCTTTCCCTACACGACGCTCTTCCGATCT), 10XP7Dual_H5 (AATGATACGGCGAC CACCGAGATCTACAC-xxxxxxxxxx-ACACTCTTTCCCTACACGACGCTCTTCCGATCT), where x represents index bases from the 10X Dual Index Kit TT, SetA H index entries. For iteration 1, the third PCR reaction was cleaned up with 0.8X SPRISelect beads and eluted in 20 µl elution buffer (EB) to generate final libraries. For iteration 2, the third PCR reaction was cleaned up with right side size selection using SPRIselect beads (0.65X beads used initially and supernatant was rebound with 1.5X beads). SABER-seq libraries were sequenced using Novaseq SP 200 cycle kits (MedGenome) with 5-10% PhiX spike-in. Sequencing parameters: Read1 28 cycles, Read2 151 cycles, Index1 10 cycles, Index2 10 cycles. Standard sequencing primers were used.

### Transcriptomic data analysis

Raw FASTQ reads were aligned against zebrafish genome, built from GRCz.109.gtf and the GRCz11.fa files, using the Cell Ranger count pipeline to generate gene-by-cell count matrices (filtered barcode matrix). The count matrices were processed using the standard workflow of the Seurat (v5.0.2) R package with modifications. Briefly, all count matrices were separately loaded into R. Seurat objects were then created with modified parameters, requiring features to be detected in at least 3 cells and cells to contain at least 300 genes. Subsequently, the Seurat objects were merged to create a single Seurat object. Cells with RNA counts between 300 and 4000 and a mitochondrial gene percentage of less than 9% were selected for downstream analysis. Gene expression values were log-normalized with the default parameters and scaled for the top 3000 variable genes. The scaled data was then used to compute the 50 principal components. Cells were iteratively clustered using the Louvain algorithm after identifying neighboring cells. Differentially expressed genes for each cluster were identified using genes that are detected in at least 10% of cells in either of the two clusters and genes showing a minimum difference of 10% between the two clusters. Cell types were annotated using known marker genes. Finally, the data was visualized using UMAP.

### SABER-seq barcode analysis

The SABER-seq FASTQ reads were processed using the Cell Ranger count pipeline, similar to the transcriptomic reads. A bash script was then used to generate a text file containing all FASTQ headers and error-corrected cell barcodes for SABER-seq data. After loading filtered barcode matrices into R, error-corrected cell barcodes from transcriptomic reads were isolated and matched against those in the text file to identify common cell barcode that are shared between the transcriptomic and SABER-seq datasets. The resulting text file contained redundant error-corrected cell barcodes associated with different UMIs for the same cell. Thus, unique error-corrected cell barcodes were isolated to create a text file containing unique FASTQ headers and error-corrected barcodes. Subsequently, FASTQ reads matching the FASTQ headers in this file were extracted from the SABER-seq raw reads using a Python script. SABER-seq barcodes were analyzed using the scGESTALT pipeline available on Github as described previously^7,35^. To analyze differences in barcode detection and editing frequencies between cell types a binomial test was used.

---

**Fig. S1.**
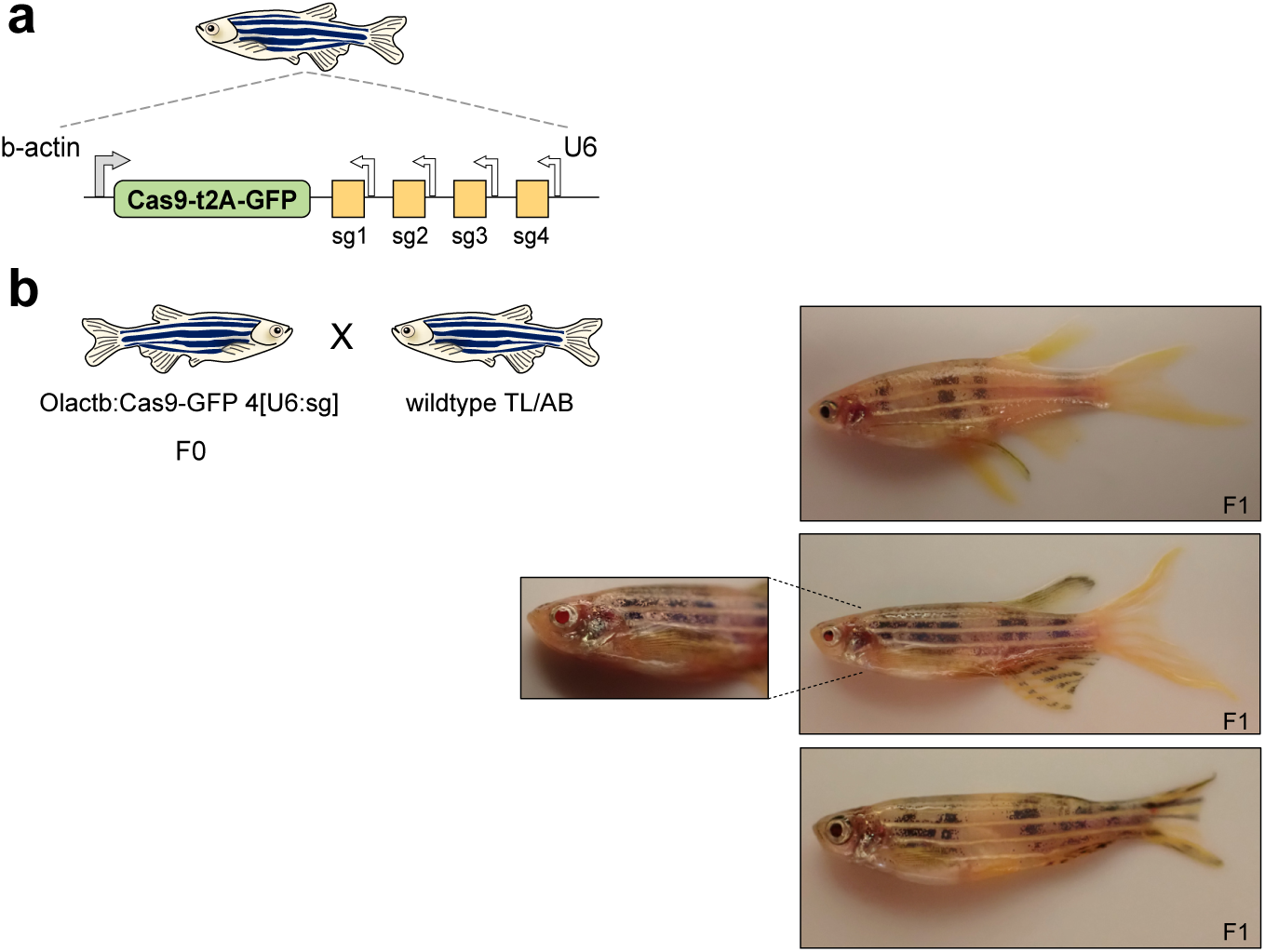
Olactb:Cas9-GFP, 4[U6:sg] transgenic adults have extensive pigmentation disruption a) Schematic of Olactb:Cas9-GFP, 4[U6:sg] transgenic reporter. sg, sgRNA. sg4 targets *tyr* gene. b) Left, schematic of Olactb:Cas9-GFP, 4[U6:sg] outcross. Right, whole-body images of adult zebrafish with mutant pigmentation in F1 adults. Dotted lines represent zoomed-in image of disruption of eye pigmentation. These transgenic animals have more extensive pigment mutations compared to TP1:Cas9-GFP, 4[U6:sg] animals due to ubiquitous expression of the b-actin promoter and hence Cas9 expression and activity.

**Fig. S2.**
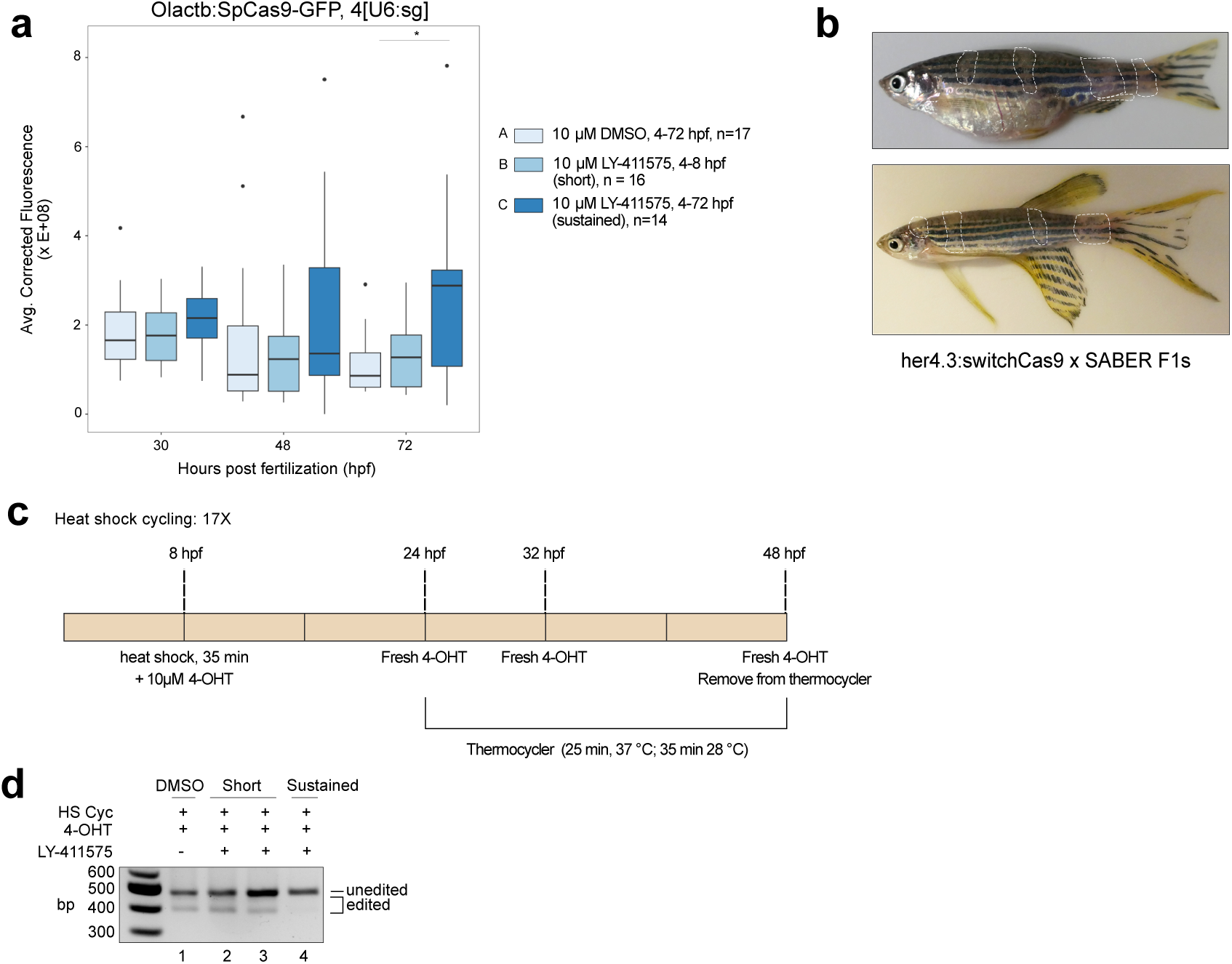
SABER, a novel CRISPR-Cas9 signal recorder. a) GFP fluorescence intensity in Olactb:Cas9-GFP, 4[U6:sg] embryos after Notch inhibition. Embryos were treated with DMSO control at 4-72 hpf, LY-411575 inhibitor at 4-8 hpf (short pulse), and LY-411575 inhibitor at 4-72 hpf (sustained). Images were taken at 30, 48 and 72 hpf. n, number of embryos. *, p < 0.05, Mann-Whitney U test. b) Whole-body images of F1 adult zebrafish from her4.3:switchCas9 x SABER cross. Dotted lines indicate areas with disrupted pigmentation in stripes. c) Schematic of heat shock cycling conditions. d) Genomic DNA amplification of SABER barcodes at 3 dpf to assess editing efficiency after Notch inhibition. HS Cyc, heat shock cycling. 4-OHT, 4-hydroxytamoxifen. LY-411575, Notch inhibitor. Embryos were treated with DMSO control at 4-72 hpf, LY-411575 at 4-8 hpf (short pulse), and LY-411575 at 4-72 hpf (sustained).

**Fig. S3.**
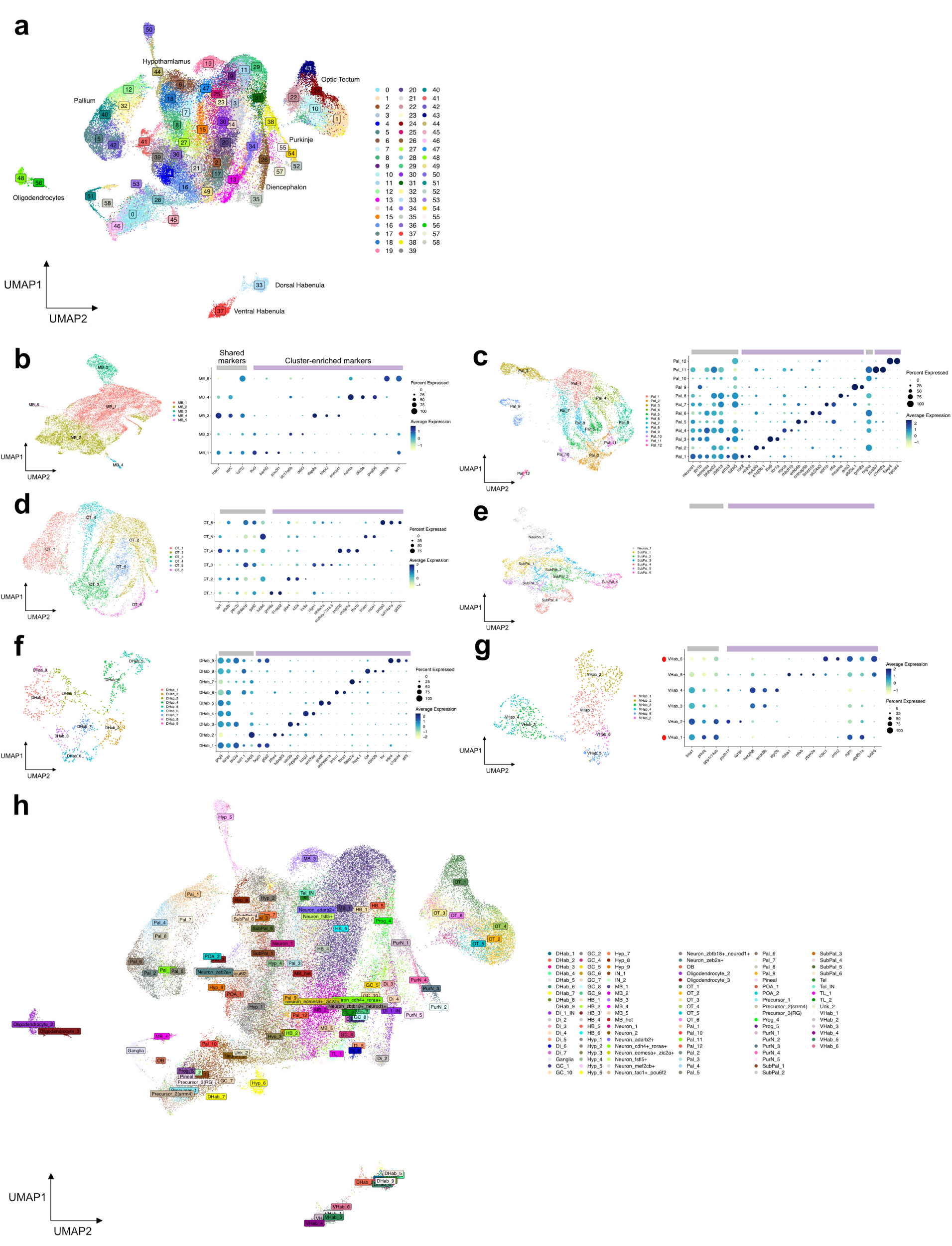
Brain cell type classification. a) UMAP embedding of 59 subclusters obtained after additional subclustering analysis of select neuron and oligodendrocyte clusters identified from the initial coarse-grained clustering shown in Fig. 4a. UMAP embedding and dot plots of marker gene expression from subclustering of b) midbrain cells, c) pallium cells, d) optic tectum cells, e) subpallium cells, f) dorsal habenula cells, and g) ventral habenula cells. Red dots represent an immature cell state (vHab_6, high tubb5 and low kiss1 expression) and a corresponding mature cell type (vHab_1, low tubb5 and high kiss1 expression). h) UMAP embedding as described in (a) with subcluster identities overlaid.

**Fig. S4.**
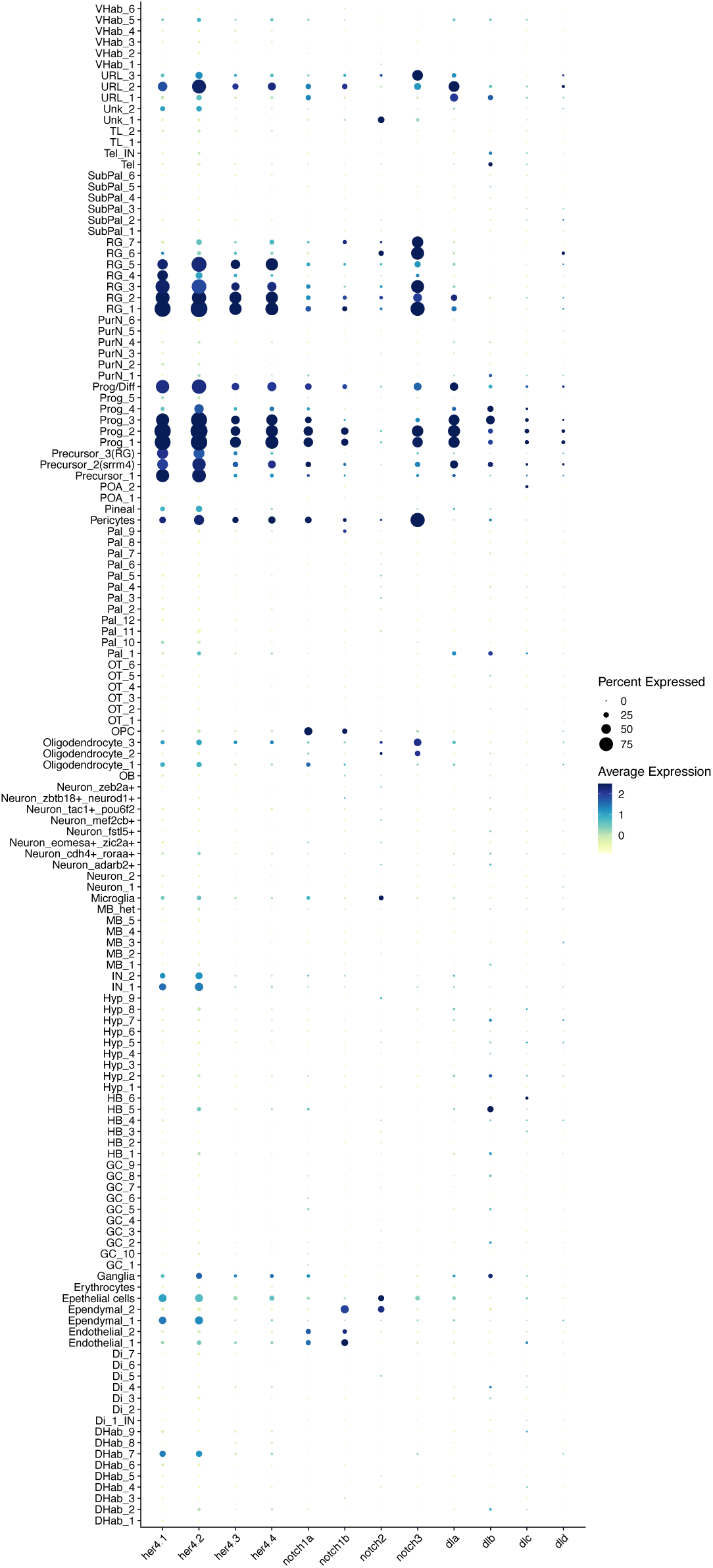
Gene expression dot plot of her4 variants, notch genes and delta genes across all subclusters.

## Notes

### Competing Interest Statement

The authors have declared no competing interest.

